# Statistical invisibility of working equids in Mexico: Dissecting the gap between global diagnostics and official data (1970–2022)

**DOI:** 10.64898/2026.04.15.718791

**Authors:** Elena García-Seco, Miguel Alonso Díaz, Tamara Tadich Gallo, Ramiro E. Toribio, Francisco Galindo Maldonado, Mariano Hernández-Gil

## Abstract

**Background:** Working equids are fundamental to the socioeconomic structure of Mexico’s small-scale agricultural sector, which accounts for 71.2% of the country’s active Agricultural Production Units (APUs). Despite their critical role in human rural livelihoods, food security, and sustainable development, these animals face systemic “statistical invisibility” within national and international productive frameworks. This study evaluates the long-term population dynamics and geographical distribution of working equids to analyze their current status amidst agricultural modernization.

**Methods:** A retrospective analysis was conducted using national census data from 1970 to 2022 provided by the National Institute of Statistics, Geography, and Informatics (INEGI). Population trends for horses, donkeys, and mules were calculated using the Average Annual Variation Rate (AAVR). The severity of population declines was classified according to an adaptation of the International Union for Conservation of Nature (IUCN) criteria. Finally, national census records from INEGI, Agri-food and Fisheries Information Service (SIAP) and The Ministry of Agriculture and Rural Development (SADER) were contrasted with FAOSTAT database estimates to identify reporting discrepancies.

**Results:** Between 1970 and 2022, the total equine population in Mexico decreased by 76.5%, falling from 6.8 to 1.6 million. However, a “paradox of modernization” was identified: while total numbers plummeted, the proportion of equids used specifically for work reached a historical peak of 81% in 2022, effectively having doubled from the 44% recorded in 2007. While donkeys and mules have suffered drastic total reductions (87% and 88%, respectively), working horses experienced a resilient 37% recovery between 2007 and 2022 (+3.71% AAVR). Furthermore, a staggering 710.8% discrepancy was found between national census data and FAOSTAT estimates, representing an overestimation of 11.3 million animals in international records.

**Conclusions:** The persistence and recent recovery of working equids reflect a “resilience of necessity” for approximately 500,000 APUs that depend exclusively on animal traction and packing due to economic constraints and complex topography. These findings challenge the narrative of total agricultural mechanization and highlight an urgent need for evidence-based public policies that address the statistical invisibility of working equids as indispensable drivers of rural sustainability and food security.

## Introduction

Small-scale agriculture remains the cornerstone of food security and rural stability in Mexico. Of the 4.6 million active agricultural productive units (APU) in the country, 71.2% are small-scale farms of five hectares or less, primarily dedicated to subsistence farming in marginalized or topographically complex terrains [1–3]. In these regions, such as the Trans-Mexican Volcanic Belt and southern highlands, modern agricultural machinery is often technically unfeasible or economically inaccessible due to the rising costs of fuel and lack of credit. Consequently, approximately 15% of all APUs (nearly 500,000 units) rely exclusively on human labor and animal traction to sustain their productive cycles [1]. In this context, working equids (horses, donkeys, and mules) are far from being symbols of technological backwardness. They represent an optimal and resilient solution that ensures rural continuity, with major economic implications [4–6]. Their role transcends tillage; they fulfill a multidimensional function by facilitating transport of water and firewood in difficult-to-access areas and connecting marginalized communities with local markets [6–10]. By mitigating the physical workload of rural families, these animals act as facilitators of human development, contributing to the “One Welfare” framework by improving the quality of life for both humans and animals [11–16].

However, despite their socio-economic relevance, working equids face significant institutional bias. Often classified as “non-productive” species in some countries, because these animals do not yield direct commodities like meat or milk, they are frequently excluded from animal health programs and national agricultural budgets [17–20]. Historically, this marginalization has been justified through a narrative of “statistical invisibility.” Nevertheless, this study argues that the primary challenge is not a lack of data, but rather data inaccuracy and a profound statistical inconsistency between global diagnostics and the official data recorded by national institutions [1–3, 21].

While international organizations often rely on “top-down” projections—frequently anchored in the stagnant dogma that 90% of global equids are primary working animals—Mexico’s INEGI has developed a “bottom-up” census methodology that meticulously tracks equine populations and their specific functions within APUs [1,11]. This disconnect between international narratives and empirical census reality has led to inaccurate welfare diagnoses and inefficient policy designs, with a negative economic impact on these communities [1–3,21]. Specifically, this has hindered the advancement of Evidence-Based Veterinary Workforce Development [22,23]. Without an accurate characterization of the targeted equine population, the veterinary educational system and public services cannot effectively strengthen the capacities needed to support these essential animals [22]; therefore, these APUs.

The present study dissects the population dynamics and geographical distribution of equids in Mexico over a 52-year period (1970–2022) [1, 24–26]. By analyzing this half-century of official data, we aim to demonstrate a “paradox of modernization“: while the total equine population may decrease due to urbanization, the proportion of animals destined for labor remains a consolidated and indispensable component of rural resilience [1,10,17,19]. Correcting the statistical gap is a fundamental prerequisite for professionalizing the educational sectors and achieving the Sustainable Development Goals (SDGs), particularly regarding poverty eradication (SDG 1) and Zero Hunger (SDG 2) [17,27,28].

## Materials and methods

### Data collection and historical scope

A retrospective analysis of working equid population dynamics in Mexico was conducted using data from the INEGI Agricultural Censuses of 1970, 1991, 2007, and 2022 [1, 24–26], spanning a 52-year period. The 1970 census was selected as the starting point because it was the first one to distinguish between the total equid population and “working equids“—specifically horses, donkeys, and mules used for traction, transport, and packing within active APUs [24]. In alignment with INEGI methodology, no distinction is made between mules and hinnies, categorizing them collectively as “mules” [1, 24–26].

Data were obtained from the Library of the Institute of Geography at the National Autonomous University of Mexico (UNAM) and the official INEGI digital platform [1,24–26]. For the 2022 period, consolidated figures were retrieved through predefined statistical tabulations to ensure accuracy [29]. These records represent the population actively integrated into agricultural systems (APUs) rather than the total national census of equids.

### Statistical Analysis and Variation Rates

Data were organized using Microsoft Excel to perform descriptive statistical comparisons across species and hybrids. To standardize the analysis of population dynamics regardless of the unequal duration between census intervals, the Average Annual Variation Rate (AAVR) was calculated [30].

A geometric growth model [31] was applied to account for the compound nature of biological population dynamics using the following formula:

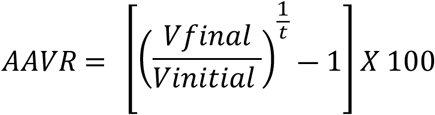

*Where V final* = population of the more recent census; *V initial* = population of the previous census, and *t* = number of years between censuses.

### Regionalization and Conservation Criteria

The national territory was delimited into five agri-food regions following The Ministry of Agriculture and Rural Development (SADER) classification [32]: Northwest (1), North (2), Central-West (3), Central (4), and South-Southeast (5) (Figure 1).

**Figure 1.**
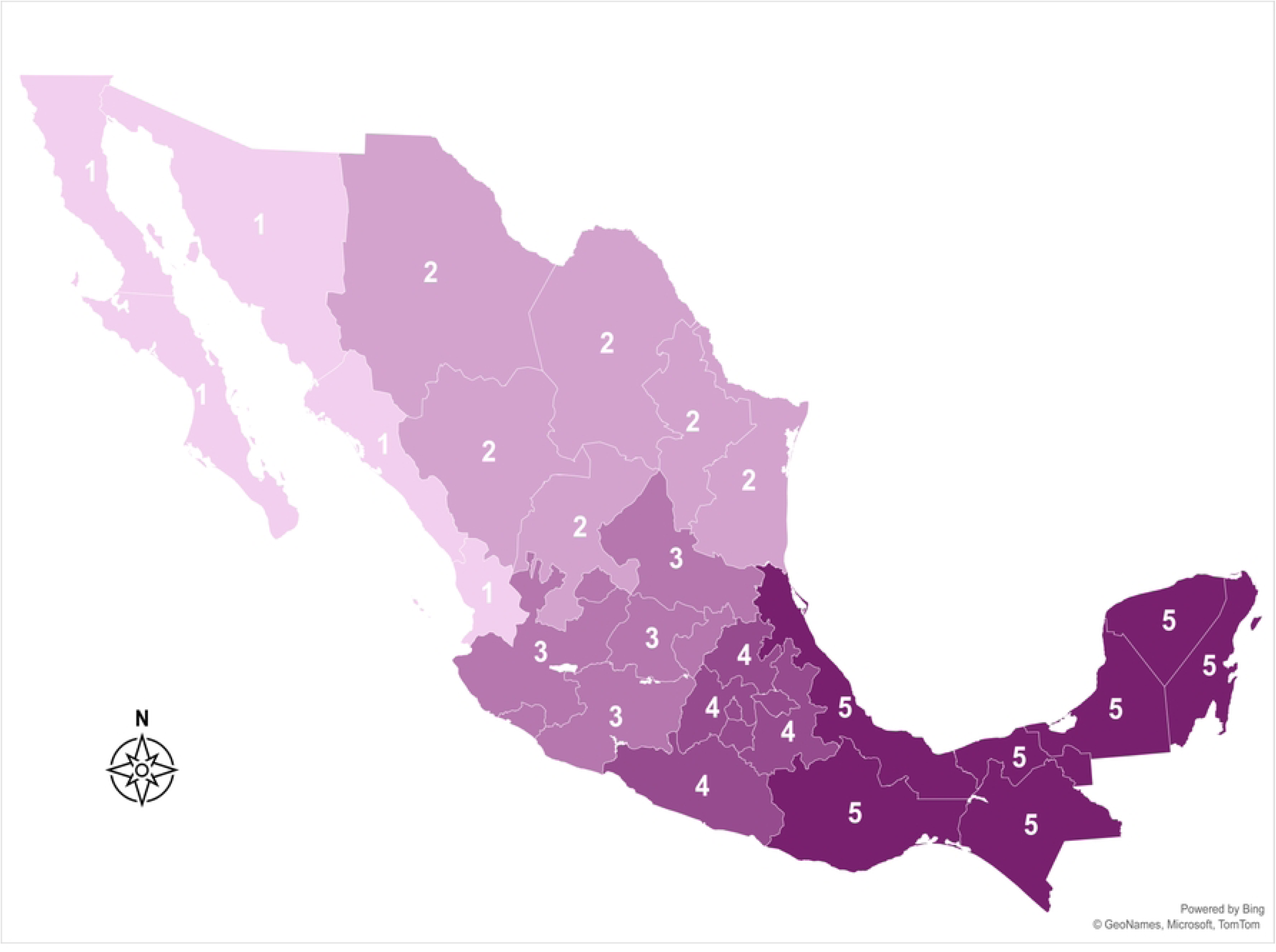
Agri-food regions of Mexico (SADER): Northwest (1), North (2), Central-West (3), Central (4), South-Southeast (5). Map generated using ArcGIS, Bing Maps, and Microsoft visualization tools based on the 2022 INEGI Agricultural Census database.

While the general population was analyzed over a 52-year period (1970-2022) [1,24–26], the distribution and population variations of working horses, donkeys, and mules were analyzed by region and federal entities using data from the two most recent censuses (2007 and 2022) [1,26]. This focus is justified by the need to assess the current conservation status of working equids. Analyzing the most recent 15-year interval allows for the application of an adaptation of the International Union for Conservation of Nature (IUCN) criteria for population reduction [33] to identify contemporary population crises and prioritize federal entities that require urgent management or conservation interventions. To evaluate the severity of these declines, federal entities were classified according to an adaptation of the International Union for Conservation of Nature (IUCN) criteria, adapted to the study’s timeframe:

- **Level 1 (Warning):** Decline 1-10%
- **Level 2 (Significant):** Decline 11-30%
- **Level 3 (Alarming/Extinction Risk):** Decline 31- 50%
- **Level 4 (Critical/Crisis):** Decline ≥ 51- 70%

### Methodological integration and validation

Geospatial distribution and map charts were generated using ArcGIS, Bing, and Microsoft visualization tools based on 2022 census data [1]. Finally, to ensure robustness and identify reporting discrepancies, INEGI census figures were compared with FAOSTAT database estimates for the year 2022 using descriptive statistics [21].

## Results

### Long-term population dynamics (1970–2022)

The total equine population in Mexico experienced a profound contraction over the 52-year study period, declining by 76.5% from 6.8 million animals in 1970 to 1.6 million in 2022. The most severe collapse occurred between 1991 and 2007, a 16-year interval during which the population plummeted by 58.6% (3 million animals) (Figure 2).

**Figure 2.**
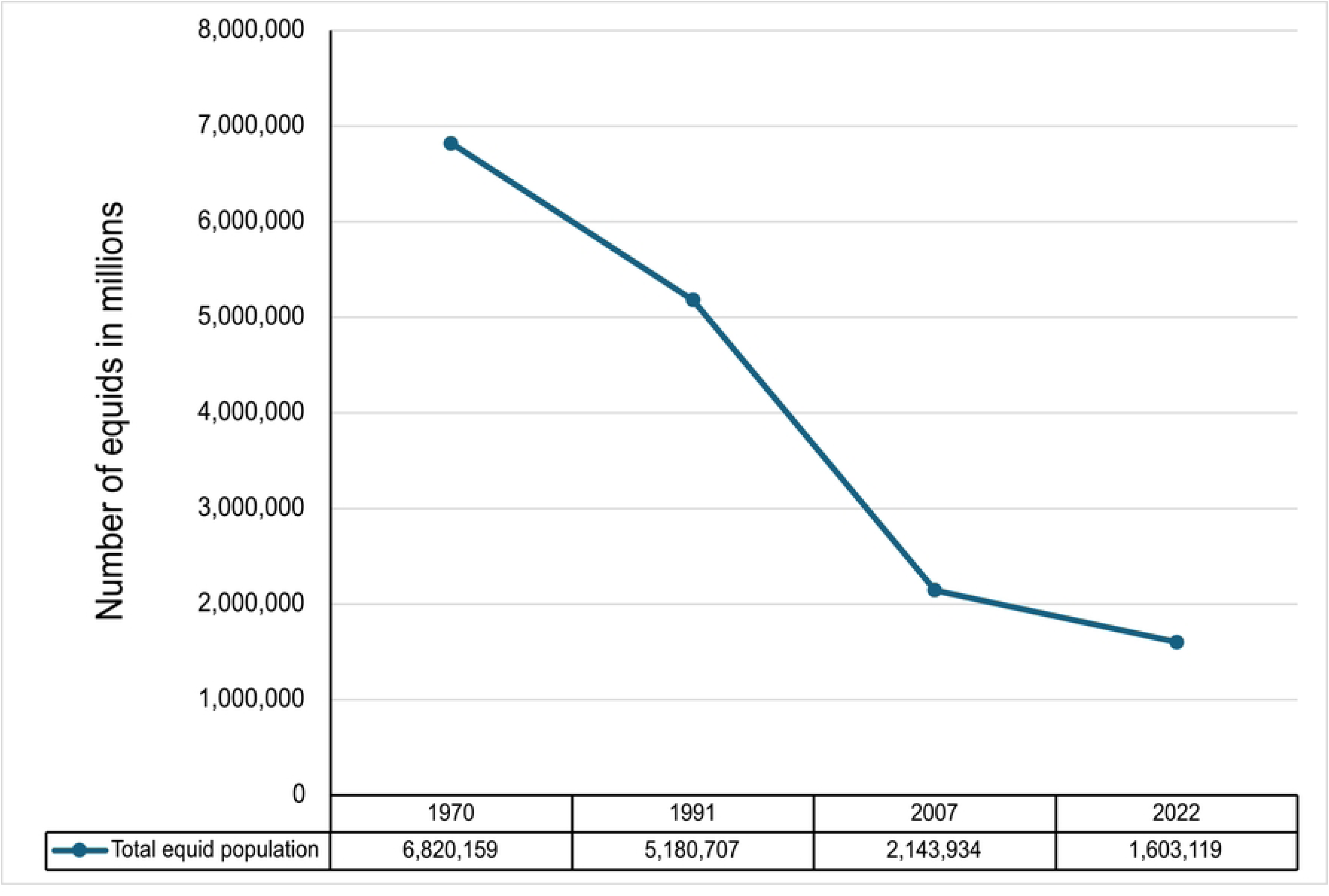
Population dynamics of working equids in Mexico (1970–2022). The graph illustrates a net reduction of 5.2 million equids over the studied period. The most significant contraction occurred between 1991 and 2007. Source: Prepared by the authors with data from INEGI Agricultural Censuses (1970, 1991, 2007, 2022).

Regarding the population specifically categorized as working equids, a parallel net reduction of 73% was recorded, falling from 4.79 million in 1970 to 1.29 million in 2022. However, a significant shift in population trends was identified in the most recent interval (2007–2022): while the total equine inventory continued to shrink, the working equid population showed a 37% recovery (Figure 3).

**Figure 3.**
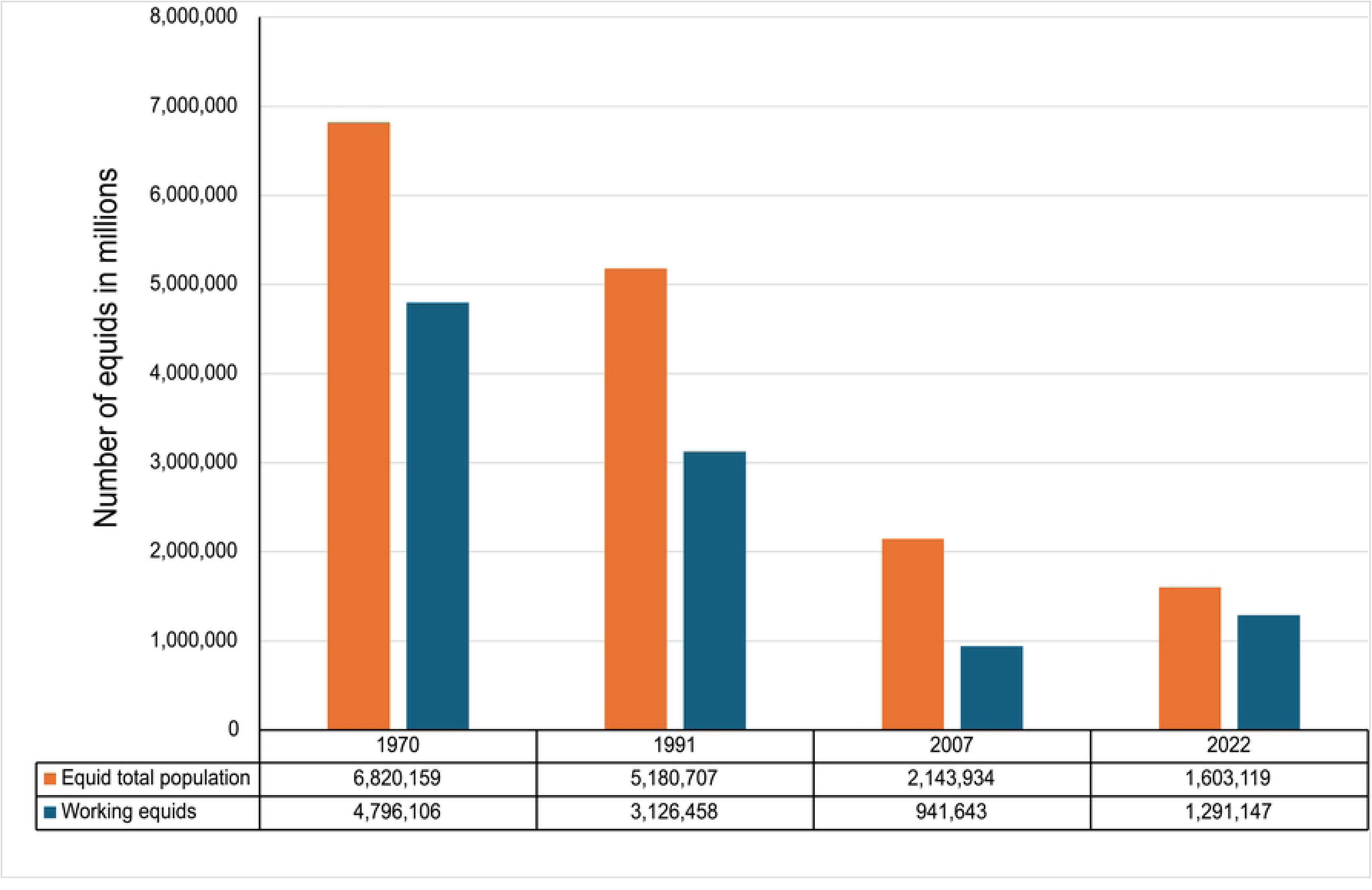
Comparative dynamics of the total equid population and working equid population in Mexico (1970-2022). The chart illustrates the parallel decline in both categories, with a significant 73% decrease in working equids over the 52-year period. The chart highlights the significant contraction of the working equid inventory, which reached its historical minimum in 2007 before a 37% recovery by 2022. Source: Prepared by the authors based on INEGI Agricultural Censuses (1970, 1991, 2007, 2022).

### The modernization paradox: utility shift and species divergence

Our analysis reveals a critical shift in the socioeconomic utility of equids in Mexico. In 1970, working animals accounted for 70% of the total equine population. This proportion reached its historical minimum in 2007 at 44%. Unexpectedly, by 2022, the ratio of working equids surged to 81% of the total inventory, effectively doubling its relative share in 15 years (Figure 4).

**Figure 4.**
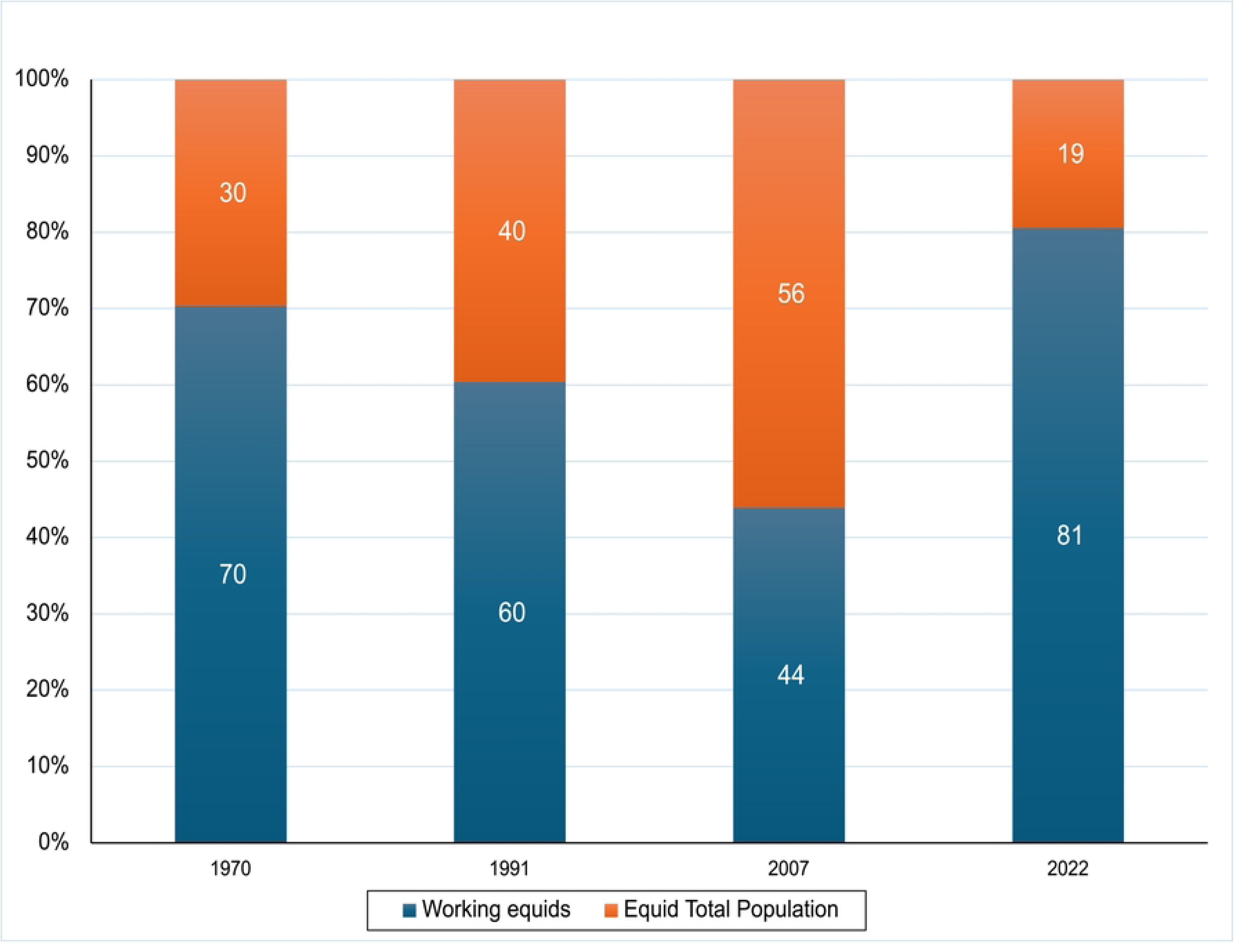
Percentage of working equids relative to the total equid population (1970-2022). The data reveals a significant shift in the utility of equids, with working animals reaching their highest relative share (81%) in the most recent census despite the overall reduction in total population size. Source: Prepared by the authors based on INEGI Agricultural Censuses (1970, 1991, 2007, and 2022).

Between the 1970 and 1991 censuses (a 21-year period), a population decline was observed across all categories. Working horses decreased by 38% (a reduction of 525,040 individuals), while donkeys and mules show a decline of 35% (a reduction of 828,774 individuals) and 30% (a reduction of 315,835 individuals), respectively. From 1991 to 2007 censuses (a 16-year period), the downward trend continued. The working horse population declined by 41% (a reduction of 353,461 individuals), donkeys declined by 80% (a reduction of 1, 216, 683 individuals), and mules experienced an 82% reduction (614,671 individuals). This resulted in a reduction in the total working equid population of 2,184,815 animals.

Finally, data from the most recent period (2007-2022) revealed a further 25% reduction (a decline of 21, 080 individuals) in the total working equid population. During these 15-year-period intervals, working horses’ population increased by 73% (an increase of 370, 584 individuals) whereas donkeys declined by 5% (a reduction of 15,740 individuals) and mules experienced 4% reduction (5,340 individuals) (Figure 5).

**Figure 5.**
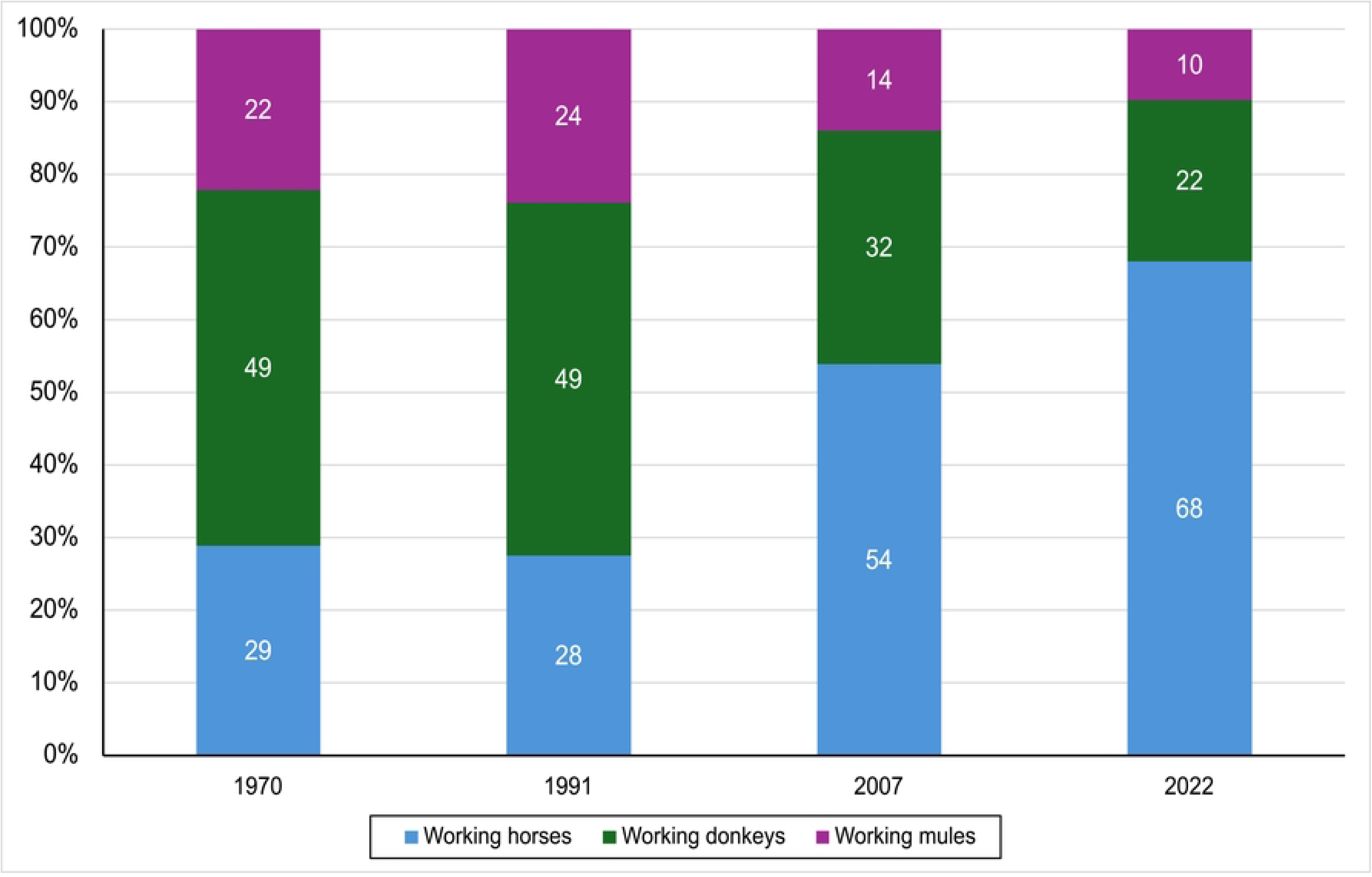
Population dynamics in percentage of working horses, donkeys, and mules (1970-2022). The stack bars represent the relative proportion (%) of each species or hybrid within the working equid population for each census year. While the total number of animals has declined since 1970, the chart highlights a significant shift in composition: working horses have become the dominant group (increasing from 29% to 68%), primarily due to the much faster rate of disappearance of donkeys and mules in recent decades. Source: Prepared by the authors based on INEGI Agricultural Censuses (1970, 1991, 2007, and 2022).

From 1970 to 2022, the working equid population underwent a drastic transformation (Fig 6). Total working horses declined by 36.6% (a reduction of 507,917 individuals), while working donkeys and mules show total reductions of 87.7% (a reduction of 2,061,197 million individuals) and 88% (a reduction of 935,846 individuals), respectively (Figure 6).

**Figure 6.**
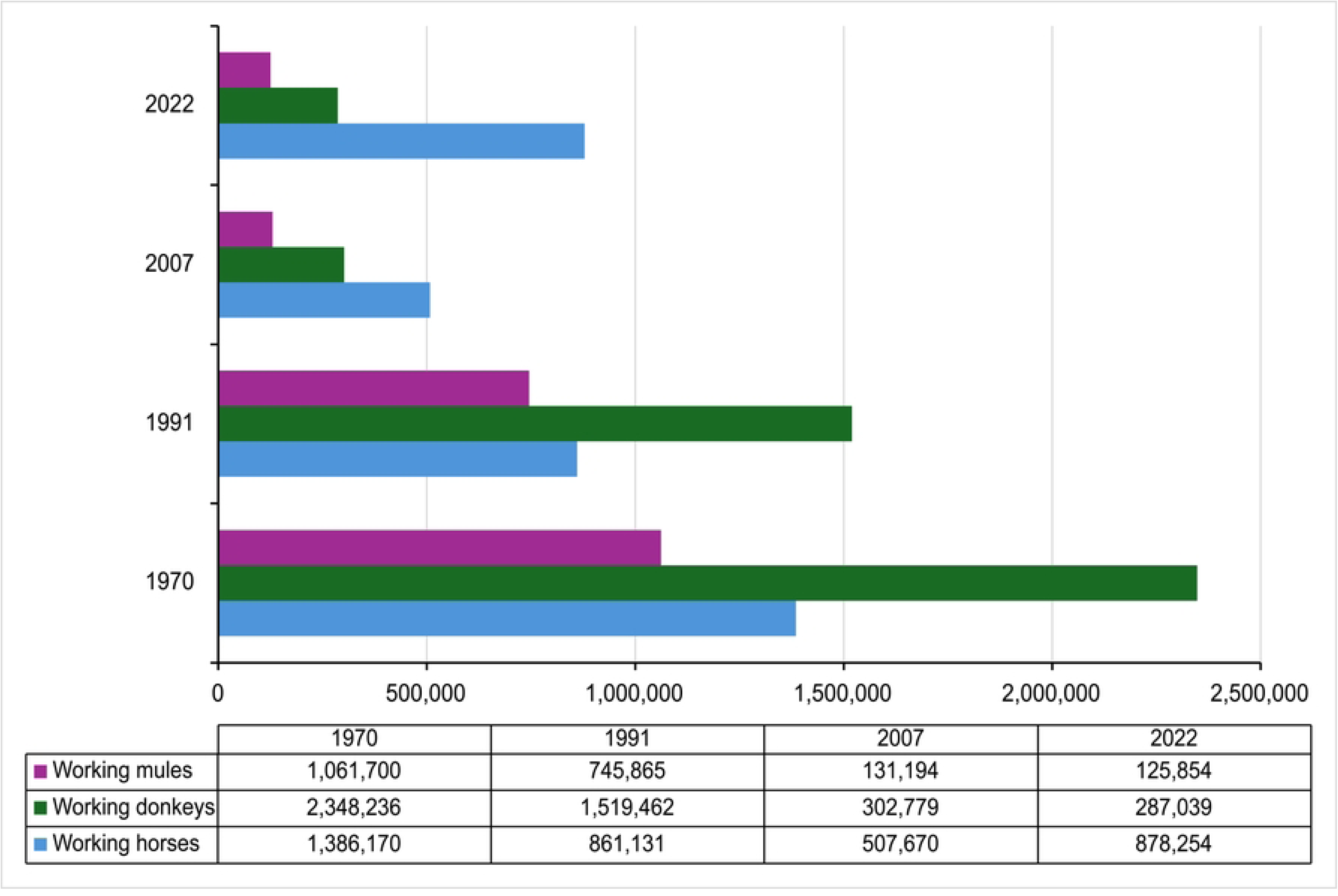
Total population of working horses, donkeys, and mules (1970-2022). The data reveals a divergent trend among species; while working donkeys and mules experienced drastic reductions of 87.7% and 88%, respectively, working horses showed a notable 37% recovery from 2007 to 2022. Source: Prepared by the authors based on INEGI Agricultural Censuses (1970, 1991, 2007, and 2022).

Prior to the detailed species analysis, it is important to highlight a significant shift in population trends. This recovery is driven by divergent dynamics among species: Working horses: After reaching a historical low in 2007, the population transitioned from a negative AAVR of −3.26% (1991–2007) to a robust recovery rate of +3.71% (2007–2022); Working donkeys and mules: These species face a much more precarious outlook. Since 1970, working donkeys have declined by 87% and mules by 88%. Although their rate of decline slowed during the 2007–2022 period (−0.35% and −0.28% AAVR, respectively), they have not shown the recovery seen in horses. While the most absolute loss in the working equid population occurred between 1991 and 2007, this 16-year interval represented the most severe contraction for all species. During this period, the population of mules plummeted from 745,865 to 131,194, while donkeys saw a drastic reduction from 1,519,462 to 302,779. Horses also reached their lowest historical AAVR at −2.60%, with their population falling from 861,131 to 507,670. This generalized decline across all species aligns with Level 4 of the IUCN criteria, signaling a critical population crisis. However, the subsequent period (2007–2022) marked a significant departure from this trend, primarily driven by the recovery of the horse population, which transitioned to a positive growth rate of +3.71%. This sharp recovery in the horse population primarily accounts for the 37% overall increase in total working equids reported in this study. The following table summarizes the population dynamics for horses, donkeys, and mules across the four evaluated census periods (Table 1).

**Table 1.** Average Annual Variation Rate (AAVR %) of equid populations across four census periods (1970–2022).

| Period | Horses (AAVR %) | Donkeys (AAVR %) | Mules (AAVR %) |
| --- | --- | --- | --- |
| 1970 - 1991 | -2.23% | -2.05% | -1.66% |
| 1991 - 2007 | -3.26% | -9.59% | -10.29% |
| 2007 - 2022 | +3.71% | -0.35% | -0.28% |

The data illustrates a significant shift in population dynamics, particularly for horses, which transitioned from a negative growth rate of −3.26% (1991–2007) to a positive recovery rate of +3.71% in the most recent period (2007–2022). In contrast, while the decline of donkeys and mules has decelerated, their growth rates remain negative, reflecting a persistent long-term reduction.

### Distribution of working equids across Federal Entities in Mexico (2007-2022)

The distribution and population variations of working horses, donkeys, and mules were analyzed by region and federal entities using data from 2007 and 2022 INEGI Agricultural Censuses are shown in Table 2 [1,26]. The Population Variations (PV) column identifies whether the population increased, decreased, or remained stable during this 15-year period. Increases and stability are indicated in parentheses, while decreases specify the percentage of reduction in the Population decline (PD) % column. Based on this last column and the IUCN criteria [33], each federal entity was classified in Level 1 (warning) to 4 (critical/crisis) (IUCN column).

**Table 2.** Regional distribution and population dynamics of working horses, donkeys, and mules in Mexico (2007–2022).

| REGION | Working Horses |  |  |  |  | Working Donkeys |  |  |  |  | Working Mules |  |  |  |  |
| --- | --- | --- | --- | --- | --- | --- | --- | --- | --- | --- | --- | --- | --- | --- | --- |
|  | Population<br>(number of animals) |  |  |  |  | Population<br>(number of animals) |  |  |  |  | Population<br>(number of animals) |  |  |  |  |
| NORTHWEST | 2007 | 2022 | PV | PD<br>% | C | 2007 | 2022 | PV | PD<br>% | C | 2007 | 2022 | PV | PD<br>% | C |
| Chihuahua | 27,436 | 43,180 | 15,744 |  |  | 7,070 | 6,805 | -265 | -4 | 1 | 5,577 | 4,455 | -1,122 | -20 | 2 |
| Coahuila | 11,828 | 24,276 | 12,448 |  |  | 4,674 | 4,966 | 292 |  |  | 1,886 | 1,206 | -680 | -36 | 3 |
| Durango | 20,842 | 40,978 | 20,136 |  |  | 7,553 | 6,856 | -697 | -9 | 1 | 8,547 | 7,416 | -1,131 | -13 | 2 |
| Nuevo Leon | 9,296 | 18,553 | 9,257 |  |  | 5,541 | 7,047 | 1,506 |  |  | 2,270 | 1,914 | -356 | -16 | 2 |
| Tamaulipas | 8,783 | 14,305 | 5,522 |  |  | 3,950 | 3,552 | -398 | -10 | 1 | 2,155 | 2,232 | 77 |  |  |
| Zacatecas | 20,712 | 41,165 | 20,453 |  |  | 9,513 | 8,363 | -1,150 | -12 | 2 | 5,977 | 3,895 | -2,082 | -35 | 3 |
| **TOTAL** | **98,897** | **182,457** | **83,560** |  |  | **38,301** | **37,589** | **-712** |  |  | **26,412** | **21,118** | **-5,294** | **-20** |  |
| **NORTH** |  |  |  |  |  |  |  |  |  |  |  |  |  |  |  |
| Baja California | 1,291 | 3,294 | 2,003 |  |  | 95 | 868 | 773 |  |  | 67 | 226 | 159 |  |  |
| Baja California Sur | 1,455 | 5,003 | 3,548 |  |  | 581 | 1,925 | 1,344 |  |  | 661 | 790 | 129 |  |  |
| Nayarit | 9,496 | 17,061 | 7,565 |  |  | 2,990 | 4,816 | 1,826 |  |  | 5,020 | 6,893 | 1,873 |  |  |
| Sinaloa | 6,225 | 12,353 | 6,128 |  |  | 2,622 | 3,367 | 745 |  |  | 3,719 | 4,039 | 320 |  |  |
| Sonora | 13,374 | 30,902 | 17,528 |  |  | 2,613 | 2,036 | -577 | -22 | 2 | 2,878 | 2,484 | -394 | -14 | 2 |
| **TOTAL** | **31,841** | **68,613** | **36,772** |  |  | **8,901** | **13,012** | **4,111** |  |  | **12,345** | **14,432** | **2,087** | **-14** |  |
| **CENTRAL- WEST** |  |  |  |  |  |  |  |  |  |  |  |  |  |  |  |
| Aguascalientes | 3,005 | 6,868 | 3,863 |  |  | 1,012 | 982 | -30 | -3 | 1 | 778 | 423 | -355 | -46 | 3 |
| Colima | 1,924 | 2,330 | 406 |  |  | 559 | 422 | -137 | -25 | 2 | 811 | 495 | -316 | -39 | 3 |
| Guanajuato | 26,847 | 32,562 | 5,715 |  |  | 16,304 | 11,764 | -4,540 | -28 | 2 | 8,453 | 5,304 | -3,149 | -37 | 3 |
| Jalisco | 18,428 | 28,355 | 9,927 |  |  | 5,639 | 4,189 | -1,450 | -26 | 2 | 7,106 | 6,298 | -808 | -11 | 2 |
| Michoacán | 24,719 | 41,017 | 16,298 |  |  | 9,109 | 8,991 | -118 | -1 | 1 | 4,953 | 5,028 | 75 |  |  |
| Queretaro | 6,802 | 12,589 | 5,787 |  |  | 4,303 | 3,789 | -514 | -12 | 2 | 1,284 | 1,242 | -42 | -3 | 1 |
| San Luis Potosi | 24,049 | 52,311 | 28,262 |  |  | 15,017 | 15,613 | 596 |  |  | 5,831 | 5,412 | -419 | -7 | 1 |
| **Total** | **105,774** | **176,032** | **70,258** |  |  | **51,943** | **45,750** | **-6,193** |  |  | **29,216** | **24,202** | **-5,014** | **-144** |  |
| **CENTRAL** |  |  |  |  |  |  |  |  |  |  |  |  |  |  |  |
| Mexico City | 1,318 | 2,003 | 685 |  |  | 137 | 129 | -8 | -6 | 1 | 248 | 235 | -13 | -5 | 1 |
| Guerrero | 40,114 | 81,271 | 41,157 |  |  | 50,741 | 61,254 | 10,513 | 21 | 2 | 17,433 | 21,600 | 4,167 |  |  |
| Hidalgo | 18,726 | 26,124 | 7,398 |  |  | 11,391 | 8,094 | -3,297 | -29 | 2 | 4,133 | 2,637 | -1,496 | -36 | 3 |
| State of Mexico | 34,666 | 56,125 | 21,459 |  |  | 24,728 | 16,835 | -7,893 | -32 | 3 | 4,177 | 3,389 | -788 | -19 | 2 |
| Morelos | 5,629 | 9,329 | 3,700 |  |  | 1,363 | 938 | -425 | -31 | 3 | 1,845 | 1,349 | -496 | -27 | 2 |
| Puebla | 34,455 | 63,018 | 28,563 |  |  | 33,588 | 29,439 | -4,149 | -12 | 2 | 12,616 | 14,027 | 1,411 |  |  |
| Tlaxcala | 5,826 | 9,379 | 3,553 |  |  | 3,632 | 2,729 | -903 | -25 | 2 | 2,981 | 2,513 | -468 | -16 | 2 |
| **Total** | **140,734** | **247,249** | **106,515** |  |  | **125,580** | **119,418** | **-6,162** |  |  | **43,433** | **45,750** | **2,317** | **-103** |  |
| **SOUTH- SOUTHEAST** |  |  |  |  |  |  |  |  |  |  |  |  |  |  |  |
| Campeche | 7,347 | 13,624 | 6,277 |  |  | 59 | 174 | 115 |  |  | 202 | 147 | -55 | -27 | 2 |
| Chiapas | 33,968 | 46,464 | 12,496 |  |  | 8,614 | 6,323 | -2,291 | -27 | 2 | 4,140 | 3,962 | -178 | -4 | 1 |
| Oaxaca | 24,977 | 37,716 | 12,739 |  |  | 40,654 | 41,881 | 1,227 |  |  | 6,388 | 7,772 | 1,384 |  |  |
| Quintana Roo | 946 | 907 | -39 | -4 | 1 | 9 | 8 | -1 | -11 | 2 | 26 | 12 | -14 | -54 | 4 |
| Tabasco | 8,649 | 22,291 | 13,642 |  |  | 39 | 109 | 70 |  |  | 246 | 186 | -60 | -24 | 2 |
| Veracruz | 52,622 | 80,642 | 28,020 |  |  | 28,655 | 22,703 | -5,952 | -21 | 2 | 8,735 | 8,232 | -503 | -6 | 1 |
| Yucatan | 1,915 | 2,259 | 344 |  |  | 24 | 72 | 48 |  |  | 51 | 41 | -10 | -20 | 2 |
| **Total** | **130,424** | **203,903** | **73,479** |  |  | **78,054** | **71,270** | **-6,784** |  |  | **19,788** | **20,352** | **564** | **3** |  |
| **Total National** | **507,670** | **878,254** | **370,584** |  |  | **302,779** | **287,039** | **-15,740** |  |  | **131,194** | **125,854** | **-5,340** | **-277** |  |

The data details the population variation (PV) and the percentage of decline (PD%) across five geographic regions, categorized by their corresponding IUCN conservation criteria level (C). Notably, while certain states show an increase in horse populations, donkeys and mules consistently exhibit percentages of decline, reaching Level 2 (Significant decline) in multiple federal entities and Level 3 (Extinction risk) in one entity.

The analysis of the last 15 years reveals a widespread decline in the working equid population across Mexico, with some federal entities falling into critical risk categories according to the adaptation of the IUCN criteria [33].

In working horses, a decrease was observed in only one federal entity at Level 1 (1-10% decline). Interestingly, 31 federal entities showed a population increase.

In working donkeys, the situation is more severe with declines in 21 federal entities. A total of 13 entities reached level 2 (11-30% decline), and 2 federal entities are at Level 3, showing a critical decline (31-50% decline). Only 11 entities show an increase. Working mules show the most widespread decline, affecting 23 federal entities. Five are at Level 1 (1-10% decline). Eleven are at Level 2, 6 have reached Level 3 (51-70%) and 1 is at Level 4 (51-70% representing a critical stage). Only 9 entities recorded an increase (Table 3).

**Table 3.** Summary of equid population trends by federal entity (2007–2022) categorized by IUCN decline levels.

| Level (IUCN) | Horses | Donkeys | Mules |
| --- | --- | --- | --- |
|  | Federal entities |  |  |
| Level 1 | 1 | 6 | 5 |
| Level 2 | 0 | 13 | 11 |
| Level 3 | 0 | 2 | 6 |
| Level 4 | 0 | 0 | 1 |
| Increase | 31 | 11 | 9 |

The data reveals a critical disparity between species; while 31 states showed an increase in working horses, 12 states have reached Level 1 (Warning), 13 states reached Level 2 (Significant decline) for donkeys and 11 states for mules. Notably, one state (Quintana Roo) has reached Level 4 (Critical) for mules, indicating a near-total collapse of specific equid populations at the regional level. (IUCN = International Union for Conservation of Nature).

### Geographical distribution of working equids (2022)

The 2022 INEGI Agricultural Census reported a total of 1,190, 226 horses with 878, 254 working horses, 287,039 working donkeys, and 125,854 working mules [1]. Population density was visualized using map charts with color scales, where darker shades indicate higher populations (Figures 7, 8, and 9).

**Figure 7.**
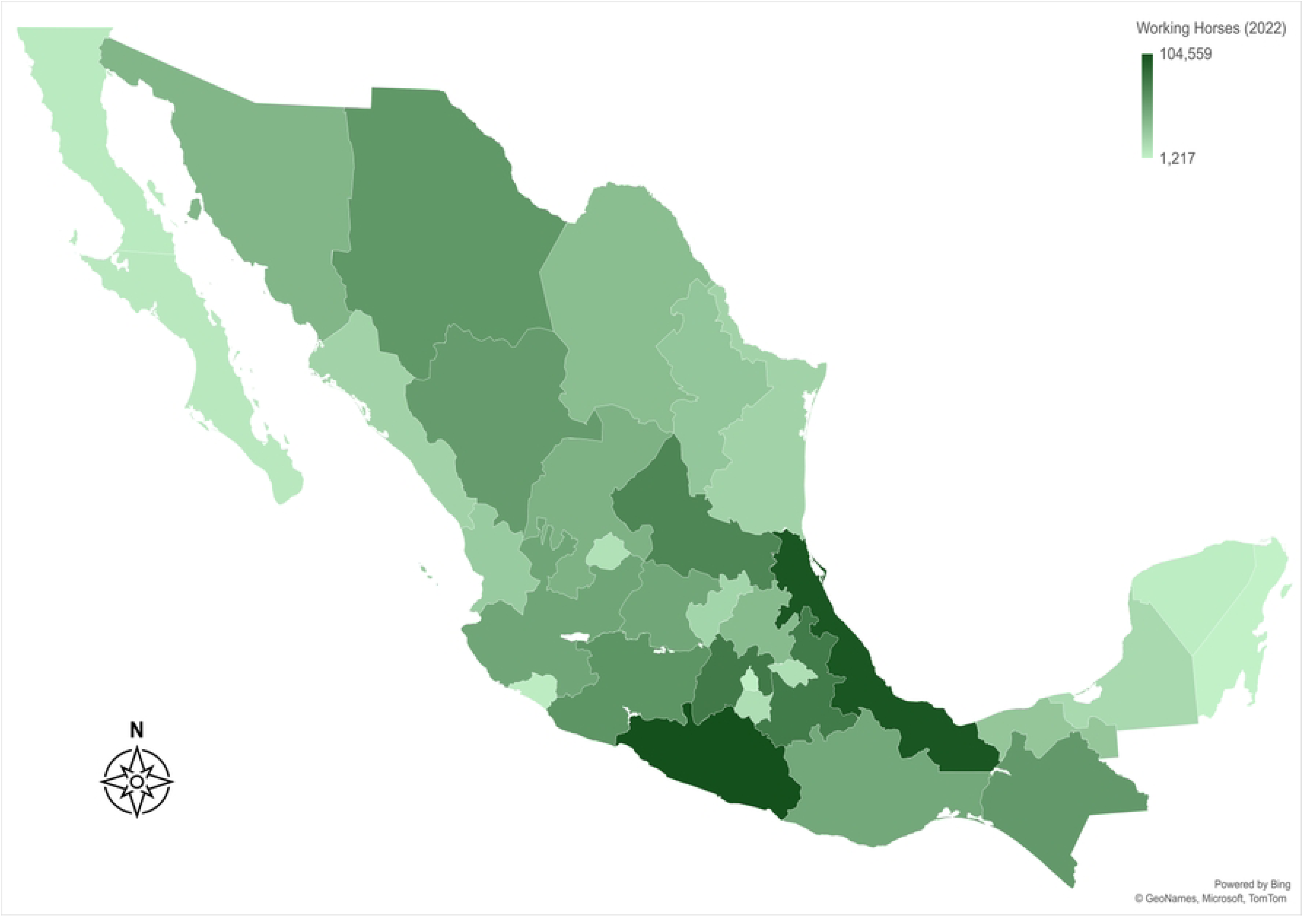
Geographic distribution of working horses in Mexico (2022). The map illustrates the distribution of horses by state, with a maximum concentration of 104,559 individuals in the most density-populated agricultural regions. The concentration of working horses is primarily in the central and southern regions of the country, reflecting their continued role in diversified agricultural systems. Map generated by the authors using ArcGIS, Bing Maps, and Microsoft visualization tools based on the 2022 INEGI Agricultural Census database.

**Figure 8.**
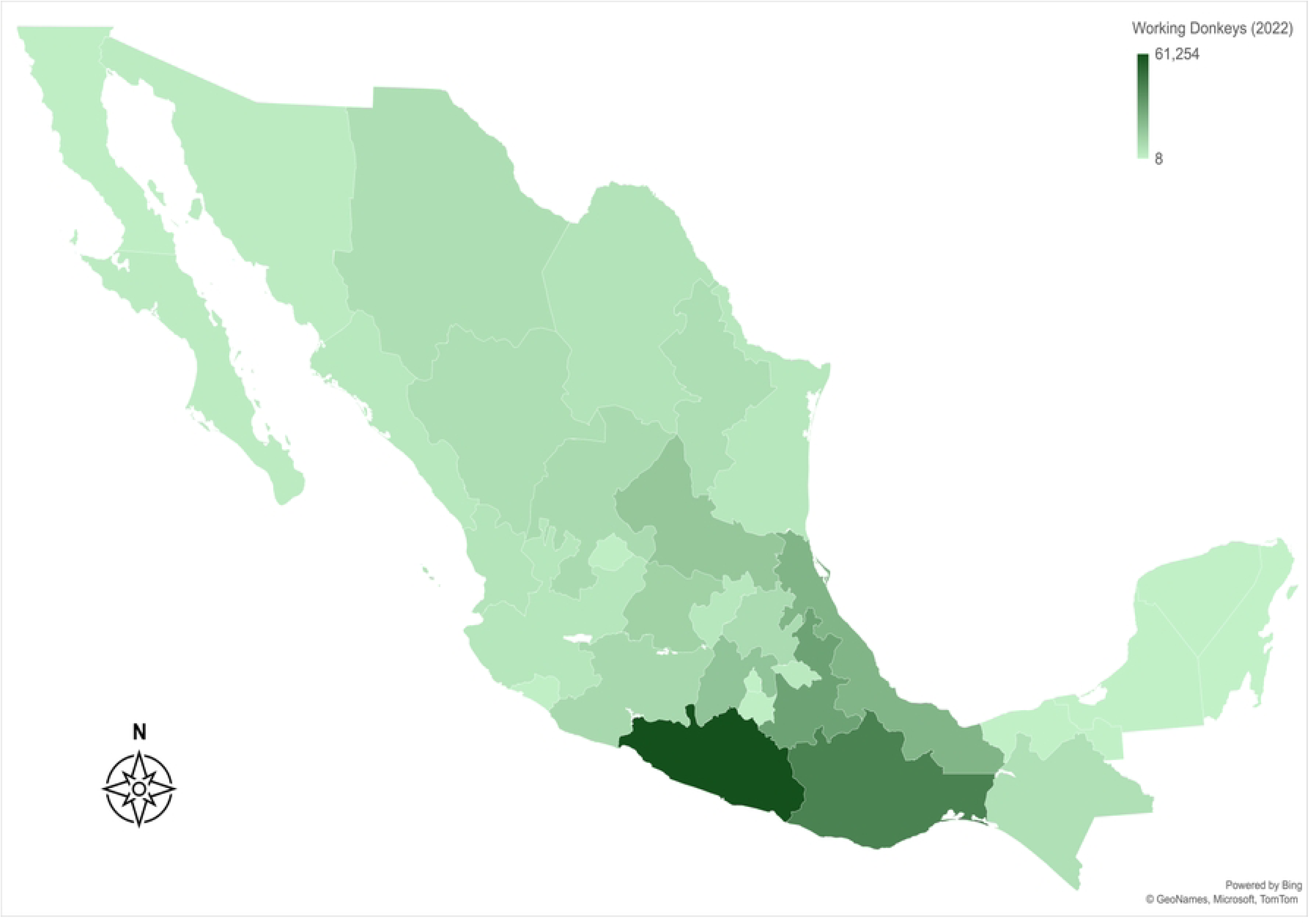
Geographic distribution of working donkeys in Mexico (2022). The map illustrates the state-level distribution of donkeys, showing a maximum concentration of 61,254 animals in the most populated states. Despite the overall national decline, specific regions maintain significant populations of working donkeys, particularly in semi-arid and mountain communities. Map generated by the authors using ArcGIS, Bing Maps, and Microsoft visualization tools based on the 2022 INEGI Agricultural Census database.

**Figure 9.**
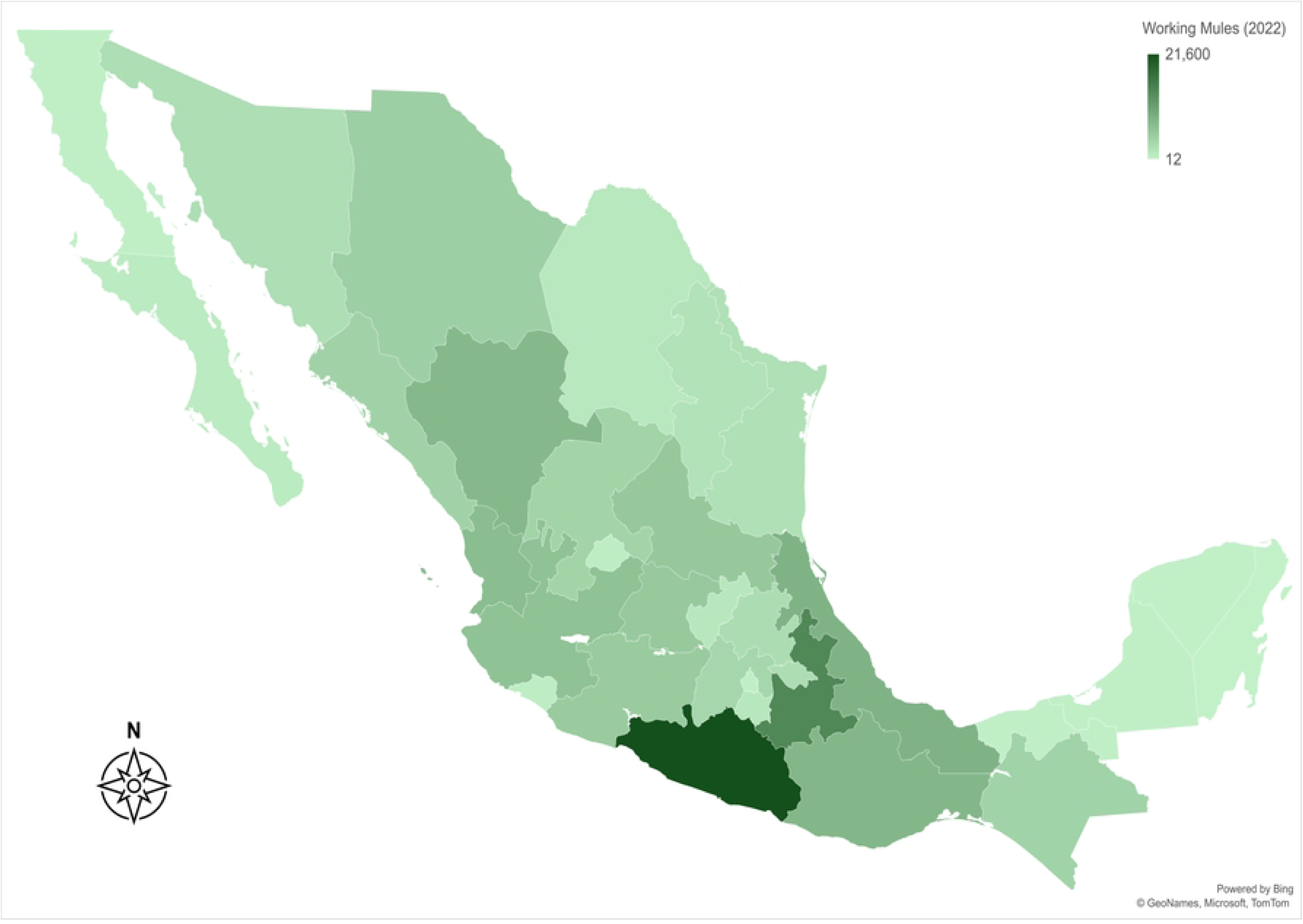
Geographic distribution of working mules in Mexico (2022). The map illustrates the density per state, with a maximum of 21,600 animals per federative entity. The data reveals that mules remain critical in specialized agricultural areas, though their distribution is more fragmented compared to horses. Map generated by the authors using ArcGIS, Bing Maps, and Microsoft visualization tools based on the 2022 INEGI Agricultural Census database.

The Central and South-Southeast regions of Mexico currently hold the largest populations of working equids. The highest concentration of horses is found in Guerrero (n = 81,217), Veracruz (n = 80,642), Puebla (n= 63,018) and the State of Mexico (n = 56,125). Donkeys are primarily concentrated in Guerrero (n= 61,254), Oaxaca (n= 41,882), and Puebla (n= 29,432), and mules are more abundant in Guerrero (n= 21,600), Puebla (n= 14,027), and Veracruz (n= 8,232). In contrast, Quintana Roo recorded the lowest population of working equids nationwide, with only 907 horses, 8 donkeys, and 12 mules [1].

### Regional distribution and IUCN vulnerability status

The distribution of working equids is highly concentrated in the Central and South-Southeast regions, particularly in states with complex topography. In 2022, the highest populations of working horses were recorded in Guerrero (n = 81,217), Veracruz (n = 80,642) and the State of Mexico (n = 77,121) [1].

Applying the adapted IUCN criteria for the 2007–2022 period reveals a widespread crisis: Mules show the most critical vulnerability, with 23 federal entities experiencing declines. Eleven states have reached Level 2 (Warning) and 6 states have reached Level 3 (Alarming/Extinction Risk) with declines of 51-70%. One state (Quintana Roo) reached Level 4 (Critical/Crisis). Donkeys are at Level 2 in 13 states and Level 3 in 2 states. In contrast, working horses showed population increases in 31 federal entities, reflecting their resilient role in specific agricultural systems.

### Statistical invisibility: FAOSTAT vs. national census discrepancy

A staggering discrepancy was identified between international estimates and national census data for the year 2022. While FAOSTAT [21] reported an estimated 12.9 million equids in Mexico, the INEGI Agricultural Census recorded only 1.6 million [1]. This represents an overestimation by international bodies of 11.3 million animals, a 710.8% difference (Table 4).

**Table 4.** Comparison of equid populations reported by FAOSTAT and INEGI in 2022. The table details the numeric difference between international estimates and national census data for horses, donkeys, and mules, showing a total discrepancy of 11,385,432 animals.

|  | FAOSTAT (2022) | INEGI (2022) | Difference between data reported by FAOSTAT and INEGI (2022) |
| --- | --- | --- | --- |
|  | Number of animals |  |  |
| Population of Horses | 6,407,072 | 1,190,226<br>(878,254 working horses) | 5,216,846 |
| **Population of Donkeys** | 3,290,520 | 287,039 | 3,003,481 |
| **Population of Mules** | 3,290,959 | 125,854 | 3,165,105 |
| **Total** | 12,988,551 | 1,603,119 | 11,385,432 |

On the other hand, Agri-food and Fisheries Information Service (SIAP) and SADER track production (meat) rather than inventory (live animals). Mexico is a top global producer of horse meat (*∼*71,000 tons/year from nearly 400,000 horses). Approximately 60-70% of horses slaughtered in Mexico (280,000 horses) are imported from the United States for “processing” as horse slaughter is prohibited there. Their meat is then exported to Europe and Japan. These animals often bypass Mexican population censuses but appear in meat production statistics [2,3]. Approximately 120,000 Mexican equids are slaughtered for the international meat market in Mexico. The transition toward mechanization (motorization) of rural areas, combined with an increasing international demand for horse meat, has likely accelerated the decline of the national equid population.

## Discussion

### The paradox of modernization and the “resilience of necessity“

The findings of this study reveal a profound paradox within Mexico’s agrarian landscape. While the total equine population has plummeted by 76% since 1970, the relative importance of working equids has reached a historical peak, accounting for 81% of the total inventory in 2022 [1]. This trend suggests that, far from being a “relic of the past,” working equids have become an essential survival strategy for small-scale producers marginalized by national mechanization policies [1–3].

The 37% recovery in the working horse population observed since 2007 does not signal rural prosperity; rather, it reflects a “resilience of necessity” driven by rising fuel costs and a lack of accessible credit for smallholders. In regions such as the Trans-Mexican Volcanic Belt, complex topography renders these animals a technical necessity where mechanical alternatives are both financially and geographically unfeasible [1].

### Statistical invisibility and data discrepancy

Working equids in Mexico face a systemic “statistical invisibility” rooted in a deep-seated institutional bias. Historically, national and international productive frameworks have classified these animals as “non-productive” species because they do not yield direct commodities such as meat, milk, or fiber [17–19]. This classification has led to their systematic exclusion from animal health programs, rural development initiatives, and national agricultural budgets. However, our findings challenge this narrative by demonstrating that working equids are not remnants of a “technological backwardness” but are, in fact, indispensable drivers of rural sustainability and food security for nearly 500,000 Agricultural Production Units (APUs) [1–3]. The failure to recognize their multi-dimensional role in transport and tillage perpetuates a cycle of neglect that hinders evidence-based veterinary workforce development and ignores their contribution to the “One Welfare” framework [11,13]. This statistical invisibility is not merely a conceptual oversight, but a documented phenomenon evidenced by the data from INEGI, SIAP, and SADER [1–3]. While international organizations like FAO often rely on broad, top-down projections—leading to the 710.8% overestimation identified in this study—the official Mexican records provide a more granular, albeit alarming, ‘bottom-up’ reality. By integrating SIAP’s figures, it becomes clear that working equids are disappearing from official productive frameworks because they do not fit the traditional ‘commodity’ model [3]. However, a profound contradiction arises at the point of slaughter. While the working equid is ‘invisible’ to national budgets and health programs, based on individual government records, approximately 400,000 animals are processed annually (yielding 70,000 tons of meat) [3,21]. This volume is comprised mainly of horses imported from the United States to be re-exported to Europe and Japan, alongside Mexican equids sold through unregulated or illegal domestic channels since there are no registered farms dedicated to breeding equids for meat in Mexico [34]. Therefore, mentioning SIAP is crucial to illustrate how national specialized agencies can expose the gaps in global diagnostics, highlighting the urgent need for a veterinary workforce that addresses this ‘invisible’ yet essential population. While statistical invisibility is a conceptual hurdle, the “data discrepancy” represents a critical technical failure in global livestock monitoring, representing an overestimation of approximately 11.3 million animals in international records [21]. Such a massive overestimation in global databases creates a false sense of security regarding equid populations, potentially masking the critical population collapses observed in donkeys and mules, leading to inefficient policy design and inaccurate welfare assessments and subsequent interventions. The data discrepancy by the FAO exists because global models assume a regulated livestock structure that does not reflect the complex, informal, and often illicit reality of the equine population in Mexico.

### Regional extinction and bio-cultural erosion

The drastic 87.7% decline in working donkeys and 88% in mules since 1970 signifies more than a demographic shift; it represents the erosion of livelihoods deeply rooted in Mexico’s rugged terrains. Analysis using the adapted IUCN criteria [33] reveals a regional crisis: while horses show resilience in 31 states, donkeys and mules have reached Level 2 and 3 in most federal entities. States such as Quintana Roo exhibit near-total collapse in their mule population. This loss of biological and cultural assets often marks the final threshold before absolute poverty for families in isolated areas.

### Policy implications and the 2030 Agenda

Recognizing working equids as fundamental drivers of food security is essential for achieving the Sustainable Development Goals (SDG 1 and 2) [27.28]. Future strategies must transition from “statistical invisibility” to institutional recognition. This includes the formal integration of equid welfare into national census frameworks and the provision of specialized veterinary services for the 500,000 APUs identified [1–3]. Only through evidence-based policies we can ensure that the families depending on these animals are no longer forced to bear the burden of rural poverty in total isolation.

## Conclusion

This study provides a comprehensive re-evaluation of the working equid population in Mexico, demonstrating that these animals remain fundamental drivers of food security and rural livelihoods. The significant 710.8% discrepancy between national census data and international reports highlights an urgent need for international organizations to rely on verified field data to avoid misinformed policies. This decoupling between international models and rural reality highlights the systemic “statistical invisibility” of the species, as international figures mask the sharp decline and the urgent need for local intervention.

The divergent population dynamics—where working horses show a resilient recovery while donkeys and mules face regional extinction—underscore the vulnerability of the peasant sector. To prevent the final collapse of these essential livelihoods, it is imperative to implement public policies that include specialized veterinary services, genetic conservation programs, and the formal integration of equid welfare into national agricultural frameworks. Recognizing working equids is essential for achieving the 2030 Agenda for Sustainable Development, particularly regarding poverty reduction (SDG 1) and Zero Hunger (SDG 2).

## Author Contribution

**Conceptualization:** Elena García-Seco, Mariano Hernández-Gil, Ramiro E. Toribio

**Data curation:** Elena García-Seco.

**Formal analysis:** Elena García-Seco, Mariano Hernández-Gil.

**Investigation:** Elena García-Seco, Mariano Hernández-Gil.

**Methodology:** Elena García-Seco.

**Supervision:** Mariano Hernández-Gil, Francisco Galindo Maldonado, Ramiro E. Toribio

**Writing-original draft:** Elena García-Seco, Mariano Hernández-Gil.

**Writing-review and editing:** Ramiro E. Toribio, Tamara Tadich Gallo, Miguel Alonso Díaz.

## Acknowledgments

UNAM Students: Luis Eduardo MolinaCorona, Javier Beltrán Romero, Erick Zuñiga Martínez, and the students from the Equine Specialty Program. This study was partially funded by the Universidad Nacional Autónoma de México (UNAM) through the DGAPA-PAPIIT project IN220424 during its first year. The funding institution had no role in study design, data collection and analysis, decision to publish, or preparation of the manuscript.

## Notes

### Competing Interest Statement

The authors have declared no competing interest.

